# Visual speech differentially modulates beta, theta, and high gamma bands in auditory cortex

**DOI:** 10.1101/2020.09.07.284455

**Authors:** Karthik Ganesan, John Plass, Adriene M. Beltz, Zhongming Liu, Marcia Grabowecky, Satoru Suzuki, William C. Stacey, Vibhangini S. Wasade, Vernon L. Towle, James X Tao, Shasha Wu, Naoum P Issa, David Brang

**Affiliations:** University of Michigan, Ann Arbor, MI 48109; Northwestern University, Evanston, IL 60208; Henry Ford Hospital, Detroit, MI 48202; Department of Neurology, Wayne State University School of Medicine, MI 48201; Department of Neurology, The University of Chicago, Chicago, IL 60637

**Keywords:** Multisensory, Audiovisual, Speech, ECoG, iEEG, sEEG, Intracranial

## Abstract

Speech perception is a central component of social communication. While principally an auditory process, accurate speech perception in everyday settings is supported by meaningful information extracted from visual cues (e.g., speech content, timing, and speaker identity). Previous research has shown that visual speech modulates activity in cortical areas subserving auditory speech perception, including the superior temporal gyrus (STG), potentially through feedback connections from the multisensory posterior superior temporal sulcus (pSTS). However, it is unknown whether visual modulation of auditory processing in the STG is a unitary phenomenon or, rather, consists of multiple temporally, spatially, or functionally distinct processes. To explore these questions, we examined neural responses to audiovisual speech measured from intracranially implanted electrodes within the temporal cortex of 21 patients undergoing clinical monitoring for epilepsy. We found that visual speech modulates auditory processes in the STG in multiple ways, eliciting temporally and spatially distinct patterns of activity that differ across theta, beta, and high-gamma frequency bands. Before speech onset, visual information increased high-gamma power in the posterior STG and suppressed beta power in mid-STG regions, suggesting crossmodal prediction of speech signals in these areas. After sound onset, visual speech decreased theta power in the middle and posterior STG, potentially reflecting a decrease in sustained feedforward auditory activity. These results are consistent with models that posit multiple distinct mechanisms supporting audiovisual speech perception and provide a crucial map for subsequent studies to identify the types of visual features that are encoded by these separate mechanisms.

## 1 Introduction

Auditory speech signals are conveyed rapidly during natural speech (3-7 syllables per second; Chandrasekaran et al., 2009), making the identification of individual speech sounds a computationally challenging task (Elliott and Theunissen, 2009). Easing the complexity of this process, audiovisual signals during face-to-face communication help predict and constrain perceptual inferences about speech sounds in both a bottom-up and top-down manner (Bernstein and Liebenthal, 2014; Lewis and Bastiaansen, 2015; Peelle and Sommers, 2015).

Multiple features extracted from visual signals can bias or enhance auditory speech perception processes, including lip shapes, rhythmic articulatory movements, and speaker identity, among others (Chandrasekaran et al., 2009; Erber, 1975; Chen and Rao, 1998; Van Wassenhove et al., 2005). While the net result is improved speech perception, each of these features may influence cortical auditory processes through distinct mechanisms. For example, visual speech is thought to influence the temporal structure of auditory speech processing by neurally amplifying auditory speech signals that are temporally correlated with lip closure, accomplished by modulating cortical excitability in auditory regions (Schroeder et al., 2008).

Indeed, functional dissociations are readily found in the auditory system. In the speech domain, research indicates that the superior temporal gyrus (STG) exhibits an anterior-posterior gradient in feature tuning, with anterior regions being more sensitive to spectral content and posterior regions being more sensitive to temporal information (e.g., broadband amplitude dynamics) (Hullet et al., 2016). Because visual speech facilitates perception for both spectral details and temporal dynamics in speech (Plass et al., 2020), it could plausibly enhance perception through multiple distinct influences on STG areas specialized for different aspects of the auditory speech signal. Importantly, prior research indicates that some audiovisual speech processes are associated with neural activity in distinct frequency bands, suggesting that they likely correspond to unique integrational functions across the sensory hierarchy (Arnal et al., 2009; Kaiser 2005; Kaiser 2006; Peelle and Sommers, 2015). Similarly, studies have demonstrated audiovisual speech effects at multiple time points, including during the observation of preparatory lip movements and after speech onset (Besle et al., 2008). However, identifying the specific role of each mechanism would be helped by first identifying different functional processes that are altered by visual speech (e.g., the modulatory effect of visual speech in different oscillatory frequency bands at different spatial and temporal scales).

Audiovisual speech integration studies using invasively implanted electrodes (intracranial electroencephalography; iEEG) have focused on raw signal amplitudes (Besle et al., 2008) or surrogate measures of population action potentials through high-gamma filtered power (HGp) (e.g., Micheli et al., 2020), showing early activation of auditory areas to audiovisual speech. However, these studies did not analyze the spectral composition of auditory-visual effects in low-and high-frequency ranges, that can reflect distinct forms of information processing (Wang, 2010; Engel and Fries, 2010; Ray, Crone, Niebur, Franaszczuk, and Hsiao, 2008), and have tended to use small sample sizes and single-participant statistics (e.g., Micheli et al., 2020; Besle et al., 2008). Conversely, non-invasive EEG studies have investigated the influence of visual speech information on low-frequency signals, with strong effects on beta and theta activity at different time scales (Sakowitz et al., 2005). However, as low- and high-frequency effects were observed across separate studies and given limitations of each approach (poor spatial resolution with EEG and small sample sizes with iEEG), the interdependence of these processes remains unclear.

Thus, at present the field lacks a unified framework for how visual speech information alters responses within auditory regions. This study sought to fill this gap by examining the interdependence of spatial, temporal, and spectral effects during audiovisual speech perception in a large cohort of patients with iEEG recordings (745 electrodes implanted in auditory areas of 21 individuals) who performed an audiovisual speech task while undergoing clinical monitoring for epilepsy. Specifically, we examined visual effects on auditory speech processes across multiple frequency bands associated with both subthreshold oscillations and neural firing. Moreover, to integrate statistical results across participants, we used linear mixed-effects models to perform statistical inference at the group level, facilitating generalization, and compared observed effects to those seen at the single participant level. Analyzing these data using group-level statistics, we found that visual speech produced multiple spatiotemporally distinct patterns of theta, beta, and high-gamma power throughout the STG. These results are consistent with the view that visual speech enhances auditory speech processes through multiple functionally distinct mechanisms and provides a map for investigating the information represented in each process.

## 2 Materials and Methods

### 2.1 Participants, implants and recordings

Data were acquired from 21 patients with intractable epilepsy undergoing clinical evaluation using iEEG. Patients ranged in age from 15-58 years (mean = 37.1, SD = 12.8) and included 10 females. iEEG was acquired from clinically implanted depth electrodes (5 mm center-to-center spacing, 2 mm diameter) and/or subdural electrodes (10 mm center-to-center spacing, 3 mm diameter): 13 patients had subdural electrodes and 17 patients had depth electrodes (Supplementary Figure 1). Across all patients, data was recorded from a total of 1367 electrodes (mean = 65, SD = 25.3, range = 24 - 131 per participant). The number, location, and type of electrodes used were based on the clinical needs of the participants. iEEG recordings were acquired at either 1000 Hz (n = 5), 1024 Hz (n = 11 participants), or 4096 Hz (n = 5 Participants) due to differences in clinical amplifiers. All participants provided informed consent under an institutional review board (IRB)-approved protocol at the University of Chicago, Rush University, University of Michigan, or Henry Ford hospital.

### 2.2 MRI and CT acquisition and processing

Preoperative T1-weighted magnetic resonance imaging (MRI) and a postoperative computed tomography (CT) scans were acquired for all participants. Registration of the pre-operative MRI to postoperative CT was performed using the ‘mutual information’ method contained in SPM12 (Viola and Wells, 1997; Penny et al., 2006); no reslicing or resampling of the CT was used. Electrode localization was performed using custom software (Brang et al., 2016; available for download online https://github.com/towle-lab/electrode-registration-app/). This algorithm identifies and segments electrodes from the CT based on intensity values and projects subdural electrodes to the dura surface using the shape of the electrode disk to counteract postoperative compression. The Freesurfer image analysis suite (http://surfer.nmr.mgh.harvard.edu/; Dale, Fischl, and Sereno 1999; Fischl, Sereno, and Dale, 1999) was used for subsequent image processing procedures including cortical surface reconstruction, volume segmentation, and anatomical labelling (http://surfer.nmr.mgh.harvard.edu/; Dale, Fischl, and Sereno 1999; Fischl, Sereno, and Dale, 1999).

### 2.3 Tasks and Stimuli

Participants were tested in the hospital at their bedside using a 15-inch MacBook Pro computer running Psychtoolbox (Kleiner et al., 2007). Auditory stimuli were presented through a pair of free-field speakers placed approximately 15 degrees to each side of the patients’ midline, adjacent to the laptop. Data were aggregated from three audiovisual speech perception paradigms (using different phonemes spoken by different individuals across tasks) to ensure generalizability of results and an adequate sample for group-analyses: 7 participants completed variant A, 8 participants variant B, and 6 participants variant C. Each task presented participants with auditory and visual speech stimuli in various combinations. As this study examines the modulatory role of visual information on auditory processes, only the auditory-alone and audiovisual (congruent auditory/visual signals) conditions were analyzed from each task variant.

On each trial a single phoneme was presented to the participant (variant A: /ba/ /da/ /ta/ /tha/, variant B: /ba/ /da/ /ga/, variant C: /ba/ /ga/ /ka/ /pa/). Figure 1 shows the timing and structure of an example trial from task variant B. Trials began with a fixation cross against a black screen that served as the intertrial interval (ITI), presented for an average of 750 ms (random jitter plus or minus 250 ms, uniformly sampled). In the audiovisual condition, the face appeared either 750 ms before sound onset (task variant B) or 500 ms before sound onset (variants A and C); across all three variants, face motion began at 500 ms before sound onset. In the auditory-alone condition, either the fixation cross persisted until sound onset (variant A) or a uniform gray square (mean contrast of the video images and equal in size) was presented for either 750 ms before sound onset (variant B) or 500 ms before sound onset (variant C). Trials were presented in a random order and phonemes were distributed uniformly across conditions. While conditions were matched in terms of trial numbers, participants completed a variable number of trials (based on task variant and the number of blocks completed): mean = 68 trials per condition (SD = 23, range = 32-96). Onset of each trial was denoted online by a voltage isolated TTL pulse.

**Figure 1:**
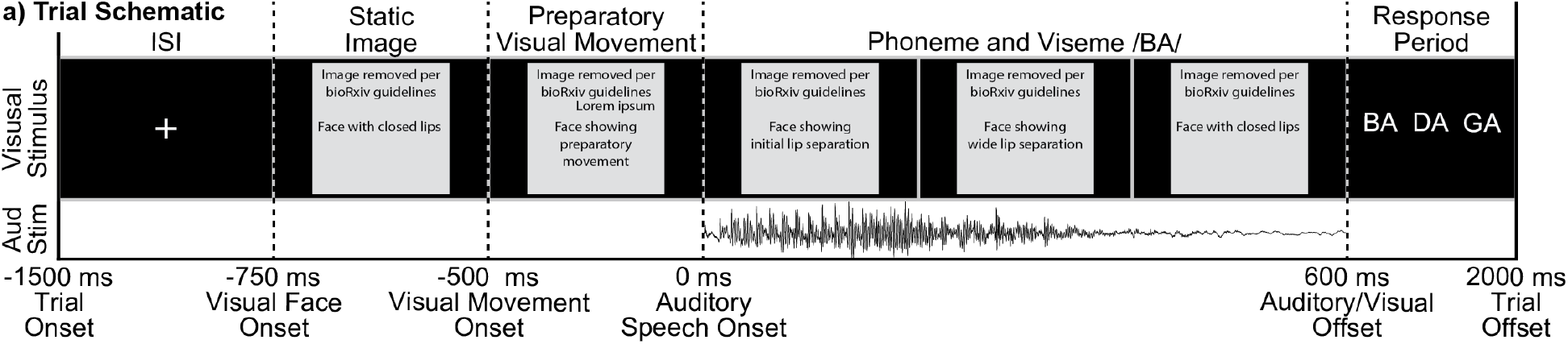
Task Variant B trial schematic. All trials began with a fixation cross 1500 ms before the onset of an auditory stimulus, lasting for an average of 750 ms (plus or minus 250 ms jitter). In the auditory-alone condition a blank screen followed the fixation cross for 750 ms. In the audiovisual condition the face appeared at the offset of the fixation (750 ms before sound onset), with preparatory visual movement beginning 250 ms later. Auditory phonemes (/ba/, /da/, or /ga/) onset at 0 ms in both conditions.

In variants A and B, following each trial participants were prompted to identify which phoneme they had heard either aloud or via button press. In variant C, participants were cued to identify a phoneme on only 20% of the trials (data not analyzed). As auditory stimuli were presented without additional noise, we anticipated high levels of accuracy. Consistent with this, in variants A and B accuracy did not differ across auditory-alone and audiovisual conditions (behavioral data was unavailable for one participant): auditory-alone mean accuracy = 95.3% (SD = 6.0%), audiovisual mean accuracy = 95.8% (SD = 6.4%), *t*(13) = 0.518, *p* = .613.

### 2.4 iEEG Data Preprocessing

Data were referenced in a bipolar fashion (signals subtracted from each immediately adjacent electrode in a pairwise manner) to ensure that the observed signals were derived from maximally local neuronal populations. Only electrodes meeting anatomical criteria within auditory areas were included in analyses. Anatomical selection required that an electrode be proximal to an auditory temporal lobe region as defined by the Freesurfer anatomical labels superiortemporal, middletemporal, and supramarginal in MNI space, resulting in 765 bipolar electrode pairs. Excessively noisy electrodes (either manually identified or due to variability in the raw signal greater than 5 SD compared to all electrodes) were removed from analyses, resulting in 745 remaining electrodes; across participants the mean proportion of channels rejected was 3.3% (SD = 8.7%, Range = 0 to 37.5%).

Slow drift artifacts and power-line interference were attenuated by high-pass filtering the data at .1 Hz and notch-filtering at 60 Hz (and its harmonics at 120, 180, and 240 Hz). Each trial was then segmented into a 2-second epoch centered around the onset of the trial. Individual trials were then separately filtered into three frequency ranges using wavelet convolution and then power transformed: theta (3 - 7 Hz, wavelet cycles varied linearly from 3-5), beta (13 - 30 Hz, wavelet cycles varied linearly from 5-10), HGp (70 - 150 Hz in 5 Hz intervals, wavelet cycles = 20 at 70 Hz, and increased linearly to maintain the same wavelet duration across frequencies); data were then resampled to 1024 Hz. Theta, beta, and HGp were selected based on previous reported findings of auditory-visual speech integration effects in these ranges (e.g., Arnal et al., 2009; Kaiser 2005; Kaiser 2006; Peelle and Sommers, 2015; Micheli et al., 2020). Within each frequency range and evaluated separately at each electrode, we identified outliers in spectral power at each time point that were 3 scaled median absolute deviations from the median trial response. Outlier values were replaced with the appropriate upper or lower threshold value using the ‘clip’ option of the Matlab command ‘filloutliers’. Across participants, a mean of .2% of values were identified as outliers (SD = .1%, Range = .1 to .5%).

Only electrodes meeting both anatomical and functional criteria were included in analyses (*n* = 745). Though electrodes were implanted in both the left and right hemispheres, electrodes were projected into the left hemisphere for visualization and analyses. This was accomplished through registering each participant’s skull-stripped brain to the cvs_avg35_inMNI152 template image through affine registration using the Freesurfer function mri_robust_register (Reuter, Rosas, Fischl, 2010). Right-hemisphere electrode coordinates were then reflected onto the left hemisphere across the sagittal axis.

Functional selection was evaluated separately for each of the three frequency bands of interest (theta, beta, and HGp) to identify auditory-responsive electrodes: accordingly, different electrodes may be included across each of the frequency analyses. To ensure orthogonality with the examined condition differences, the functional localizer required electrodes to demonstrate a significant post-stimulus response (0 - 500 ms) regardless of condition relative to zero using one-sample t-tests after correcting for multiple comparisons using false discovery rate (FDR). Beta and theta selection applied two-tailed t-tests while HGp applied one-tailed t-tests (as meaningful auditory HGp responses were predicted to elicit HGp increases (Beauchamp., 2016)).

### 2.5 Group-Level Analyses

Traditionally, iEEG studies have focused on individual-participant analyses utilizing fixed-effect statistics (e.g., Micheli et al., 2018; Besle et al., 2008; Chang et al, 2010; Plass et al., 2020). While these approaches are valid for estimating parameters and effect sizes within a single individual, they do not provide estimates across participants and thus lack generalizability across epilepsy patients, making inferences to the general population more difficult. Moreover, some studies mix between- and within-participant statistics by aggregating data from all participants without modeling participant as a random effect, violating independence assumptions (e.g., Lega, Germi, Rugg, 2017). This approach has been discussed extensively under the title of ‘pseudoreplication’ and can lead to spurious and poorly generalized results (for a discussion see Aarts et al., 2014; Lazic 2010; Lazic et al., 2018). These concerns for iEEG research have been raised and theoretically addressed previously by other groups using variants of a mixed-effects model (Kadipasoglu et.al., 2014; Kadipasoglu et al., 2015). To overcome these limitations, we employed two separate analysis approaches.

#### 2.5.1 Group-Level Spatial Analyses

To identify regions of the auditory temporal lobe that responded differently to auditory-alone versus audiovisual stimuli, we conducted individual-participant statistics and aggregated data across participants using an approach from the meta-analysis literature (treating each participant as an independent replication). Specifically, each ‘virtual’ bipolar electrode (calculated as the average coordinates between the associated pair of electrodes) was transformed into MNI space (Freesurfer cvs_avg35_inMNI152) and linked to neighboring vertices (within 10 mm Euclidean distance) on the Freesurfer MNI cortical pial surface (decimated from 1 mm to 4 mm); this one-to-many approach mitigates the imperfection of cross-participant spatial registration. Next, statistics were evaluated separately at each vertex for each participant using independent-sample t-tests, to compare auditory-alone and audiovisual trials between −1000 to 500 ms (auditory-onset at 0 ms; data were averaged across 100 ms time-windows prior to statistical analyses). Within-participant statistics were adjusted for multiple comparisons across vertices and time using FDR (Groppe, Urbach, and Kutas, 2011). The approach yielded individual-participant p-value maps at each of the 15 time-points. P-value maps were then aggregated across participants using Stouffer’s Z-score method (Stouffer et al., 1949).

#### 2.5.2 Group-Level Regional Time-Series Analyses

While the meta-analysis approach establishes the strength of an effect at the group-level, it fails to provide group-level estimates and cannot effectively model data from both within and between participants (as is necessary in the evaluation of interactions across time, space, and analyzed frequency ranges). To model more general group-level differences between auditory-alone and audiovisual conditions we used linear mixed-effects models. Because appropriately fitted models require more data than is often present at a single vertex, we created three regions of interest (ROIs) within the STG. ROIs were divided into three equal partitions from the “superiortemporal” label in Freesurfer, comprising anterior, middle, and posterior regions, similar to the division of the STG used previously (Smith et al., 2013). Electrodes within 10 mm of these labels were linked to the closest of the three (no electrode was linked to multiple labels). Our focus on the STG was motivated by previous demonstrations of strong effects of lipreading in this region (e.g., Smith et al., 2013). A numerical breakdown of the number of electrodes and participants in each of the three regions of the STG is provided in Table 1.

**Table 1:** Number of electrodes and participants present in each of the three regions of STG analyzed.

|  | **Anterior STG** |  | **Medial STG** |  | **Posterior STG** |  |
| --- | --- | --- | --- | --- | --- | --- |
| **Frequency** | **N. Elecs** | **N Partic.** | **N. Elecs** | **N Partic.** | **N. Elecs** | **N Partic.** |
| **HGp** | 22 | 10 | 150 | 18 | 66 | 12 |
| **Beta** | 59 | 14 | 138 | 18 | 62 | 12 |
| **Theta** | 72 | 16 | 162 | 18 | 72 | 12 |

Linear mixed-effects modeling was performed using the fitlme function in Matlab R2019a (Mathworks Inc., Natwick, MA). Electrodes in the same ROI from the same participant were averaged prior to analysis to reduce the complexity of the model and as neighboring electrodes share variance. Individual trials were not averaged within or across participants prior to analysis. Nine main-effect models were constructed, in which differences between auditory-alone and audiovisual trials were separately evaluated at each of the three STG ROIs (anterior, middle, posterior) and three frequency bands (theta, beta, HGp) using the equation:

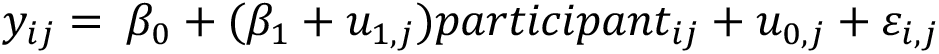

Where, *y* represents the ECoG trial value, with a fixed effects term for the trial condition and a random intercept and slope term for the participant ID. In Matlab notation, this is represented as: ECoG_Trial_Value ~ Trial_Cond + (Trial_Cond|Participant_ID). Critically, we modeled both random intercepts and random slopes for trial condition as there were multiple measurements per participant and to maintain ‘maximal’ models for confirmatory hypothesis testing (Barr et al., 2013). Statistics for the main-effect models were adjusted for comparisons at multiple time-points from −500 to 500 ms using FDR correction (*q* = .05) (Groppe, Urbach, and Kutas, 2011).

Interaction models were subsequently constructed to evaluate whether audiovisual versus auditory-alone condition effects varied as a function of frequency band, ROI, and time, using the Matlab notation: ECoG_Trial_Value ~ Trial_Cond * FrequencyBand * ROI * Time + (Trial_Cond|Participant_ID). While these model parameters were selected for inclusion based on confirmatory hypothesis testing, we also justified model selection using AIC comparisons. Separate models were constructed at each 1 ms time-point for the main-effect models (shown in Figures 6–8). Data were averaged in 5 ms time-bins for the interaction models due to computational complexity and memory requirements. The inclusion of time as a random factor in interaction models may appear to violate the assumption of independence as spectral power demonstrates autocorrelations. However, the inherent characteristics of the mixed effect model’s covariance structure should account for this dependence (Riha et al., 2020; Barr et al., 2013). More generally, calculating degrees of freedom with linear mixed-effect models is a readily acknowledged challenge (e.g., Luke 2017). Acknowledging this, model significance was estimated using residual degrees of freedom. To ensure that the likely inflated degrees of freedom did not drive our effects, we additionally examined effects using a conservative estimation of degrees of freedom, based only on the number of participants who contributed data to a particular analysis (maximum of 21); all interactions that were significant remained significant (at p<.001).

### 2.6 Individual Electrode Analyses

To examine individual differences in the patterns of activity evoked across electrodes and participants, individual electrode statistics were examined at representative electrodes. Unpaired t-tests were conducted separately at each time-point comparing audiovisual versus auditory HGp (random factor = trial). Statistics for the main-effect models were adjusted for comparisons at multiple time-points from −500 to 500 ms using FDR correction (*q* = .05).

## 3 Results

### 3.1 Group-Level Spatial Analyses

Figure 2 shows the spectro-temporal plot of the event related spectral power (ERSP) for audiovisual signals from all auditory electrodes across all participants. Data demonstrate that spectral power was distributed over multiple frequency bands while audiovisual stimuli were presented: increased power in theta and high-gamma ranges, along with beta suppression. This, supported by past studies, provides justification for subsequent analyses focusing on these three frequency bands.

**Figure 2:**
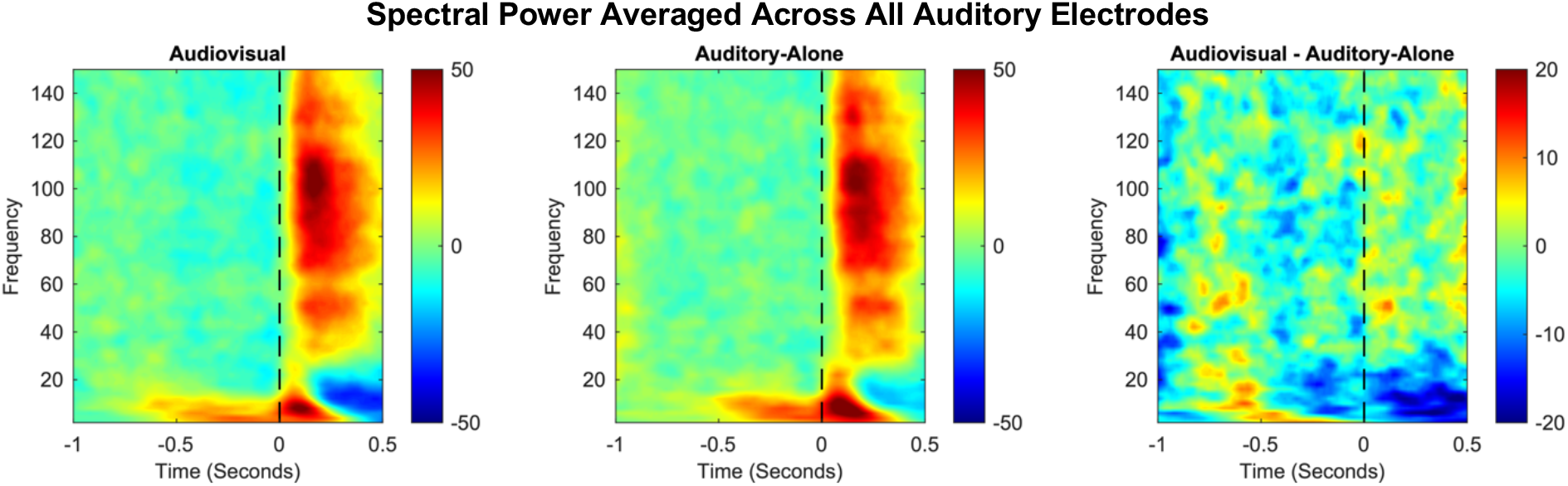
Group-level plots showing event-related spectral power from 2-150 Hz. Data reflect ECoG activity from all anatomically localized electrodes (*n* = 745), first averaged across electrodes within each participant, then averaged across participants. Dotted lines denote auditory onset. Color scale reflects normalized power.

### 3.2 Group-Level Spatial Analyses: Theta Power

Figure 3 shows group-level differences in theta power (3 - 7 Hz) between audiovisual and auditory-alone trials. A small but significant difference (audiovisual > auditory) emerged from −700 to −600 ms before sound onset in the supramarginal gyrus (peak coordinates: x = −60.7, y = −56.2, z = 30.3, *p* = 0.001) with a peak-response in this region between −600 to −500 ms before sound onset (peak coordinates: x = −60.7, y = −56.2, z = 30.3, *p* = 0.0003). This activation pattern reflected only a small percentage of the supramarginal gyrus (SMG) (1.7% of SMG vertices at time-point -700 to −600 ms, and 2.6% of SMG vertices at time-point −600 to −500 ms). In contrast to this initial pattern, the majority of condition differences were observed in the middle temporal gyrus (MTG) and STG with significantly more power in auditory trials compared to audiovisual trials. This pattern emerged as early as −300 to −200 ms (peak coord: x = −47.2, y = −33, z = −4.3, p = 0.0003) and peaked during the time-range 100 to 200 ms following sound onset (peak coord: x = −60.8, y = −20.5, z = 11.4, p = 9.6e-12). The greatest proportion of significant vertices were observed from 200 to 300 ms (STG = 27.5%, MTG = 12.4%, SMG = 5.4%), strongly weighted towards the middle to posterior STG. These data suggest that the majority of theta-related activity during audiovisual speech processing occurs following sound onset.

**Figure 3:**
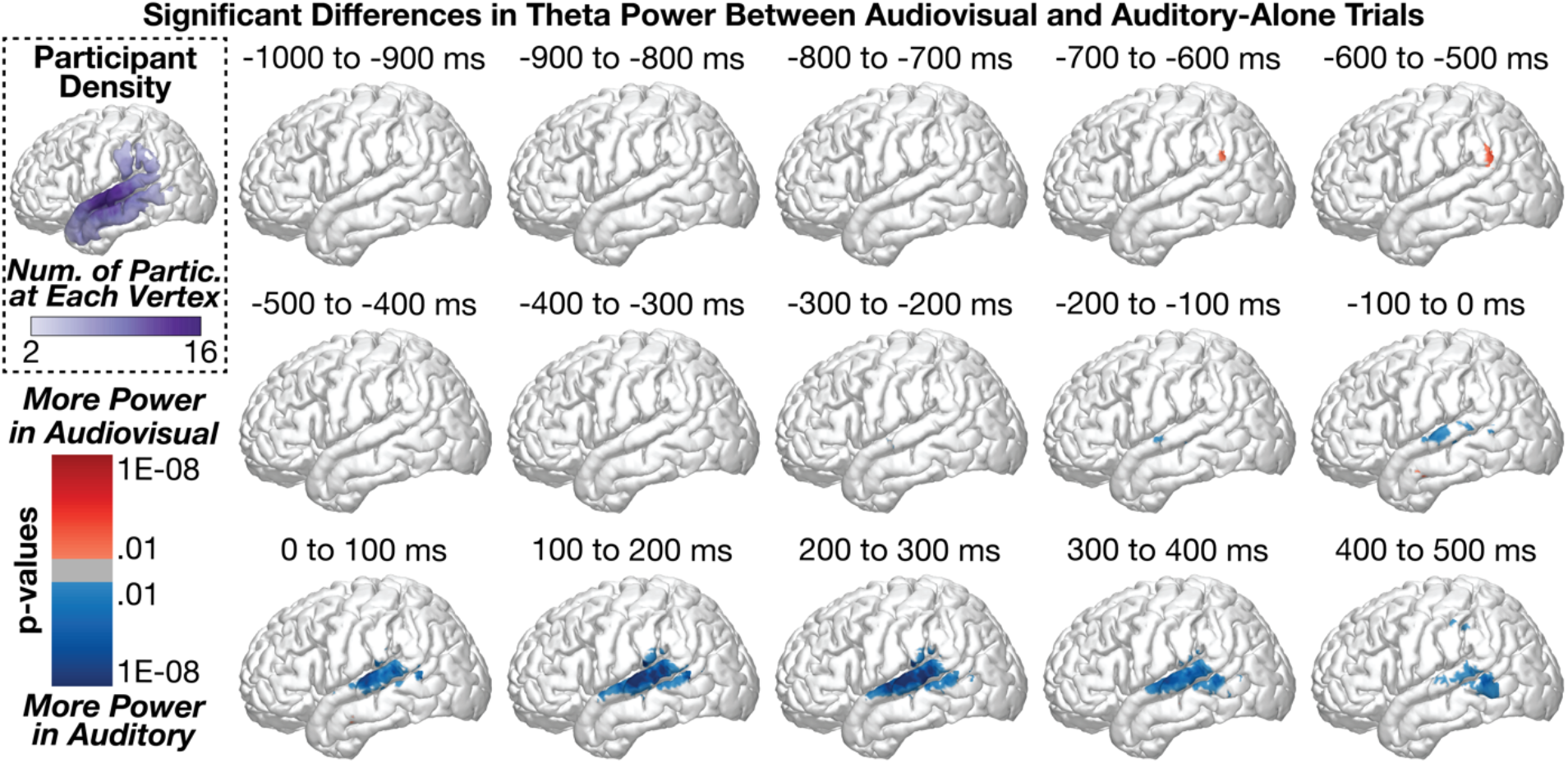
Group-level analyses comparing theta power between audiovisual and auditory-alone conditions at 100 ms time-windows (sound onset at 0 ms). Statistics conducted vertex-wise at the individual participant level and aggregated across participants using Stouffer’s Z-Score method. Multiple comparisons applied across time and space using FDR. Top-left plot shows the number of participants who were included at each vertex. Congruent audiovisual stimuli elicited reduced theta power at the middle to posterior STG, peaking after the onset of the speech sound.

### 3.3 Group-Level Spatial Analyses: Beta Power

Figure 4 shows group-level differences in beta power (13 - 30 Hz) between audiovisual and auditory-alone trials. As was observed in the theta band, a small but significant difference (audiovisual > auditory) emerged from −700 to −600 ms before sound onset in the supramarginal gyrus (peak coordinates: x = −60.1, y = −24.6, z = 15, *p* = 0.005; .2% of SMG vertices were significant); no other significant audiovisual > auditory differences were observed throughout the time-series. In contrast to this initial pattern, the majority of condition differences were observed in the STG with significantly more power in auditory trials compared to audiovisual trials; this observation of reduced beta power is most consistent with *increased* beta suppression (see Section 3.7 for additional evidence). This pattern emerged as early as −400 to −300 ms (peak coord: x = −61.8, y = −1, z = −11.8, *p* = 0.0001) along the anterior to middle STG/MTG and peaked −200 to −100 ms before sound onset (x = −65, y = −10, z = 0.9, *p* = 8.8e-08); the majority of significant vertices during this time-range were in the STG: STG = 16.2%, MTG = 3.1%, SMG = 2.7%. Whereas the peak activation occurred from the −200 to −100 ms time-window, the greatest proportion of significant vertices were observed in the −100 to 0 ms time-window range: STG = 20.6%, MTG = 4.9%, SMG = 2.3%. These data suggest that the majority of beta-related activity during audiovisual speech processing occurs before sound onset in contrast to the spatial and temporal pattern of results observed for theta band activity. See Section 3.9 for a direct comparison of the spatiotemporal effects between theta and beta band activity. As the differences did not emerge until after face-onset but immediately prior to sound onset (i.e., during which time preparatory visual movements were observed by participants), we interpret these results to reflect predictive coding information along the STG (e.g., Bastos et al., 2012; Peelle and Sommers, 2015).

**Figure 4:**
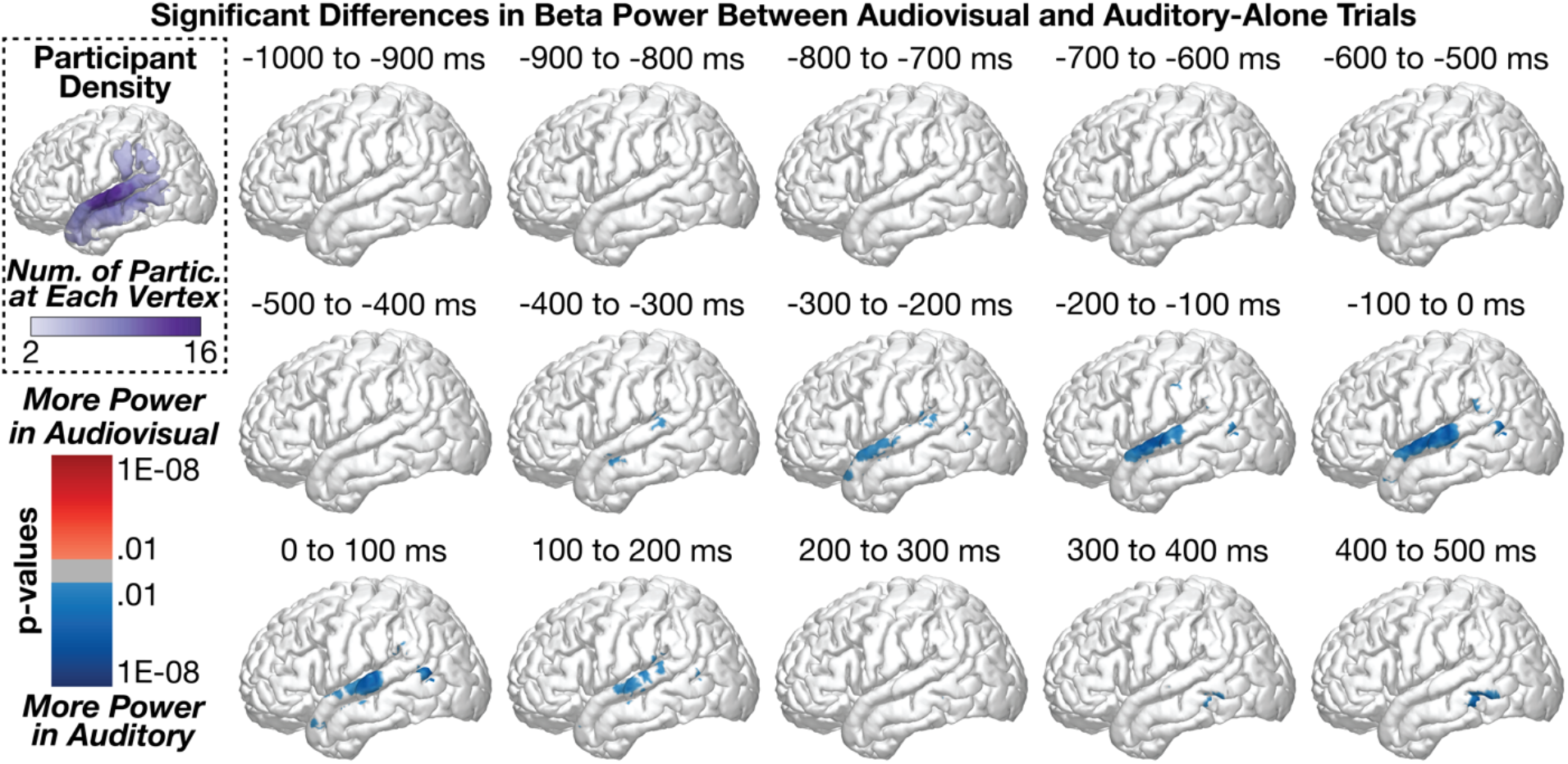
Group-level analyses comparing beta power between audiovisual and auditory-alone conditions at 100 ms time-windows (sound onset a 0 ms). Top-left plot shows the number of participants who contributed data to each vertex. audiovisual stimuli elicited greater beta suppression at the posterior STG, peaking before sound onset.

### 3.4 Group-Level Spatial Analyses: High-Gamma Power

Figure 5 shows group-level differences in high-gamma power (HGp; 70 - 150 Hz) between audiovisual and auditory-alone trials. The first significant time-points in the series were observed in the MTG (audiovisual > auditory) beginning from −700 to −600 ms (peak coord: x = −55.8, y = - 63.2, z = 8.4, *p* = 1.5e-07). Small clusters of effects were observed between −600 to −100 ms (all effects reflected less than 5% of the number of vertices in each region). Beginning from −100 to 0 ms, however, we observed a strong cluster of significant differences (audiovisual > auditory) in the MTG and STG (peak coord: x = −57.4, y = −66.6, z = 9.4, *p* = 8.1e-12, Region = MTG, percent significant vertices in each region: STG = 8.2%, MTG = 10.1%, SMG = 1.7%). This effect persisted throughout the time-series but shifted more inferior to the MTG by 400 to 500 ms (proportion significant vertices in each region: STG = 1.4%, MTG = 12.2%, SMG = 0%). In contrast to results in the theta and beta frequency bands, HGp effects were largely restricted to the posterior STG/MTG.

**Figure 5:**
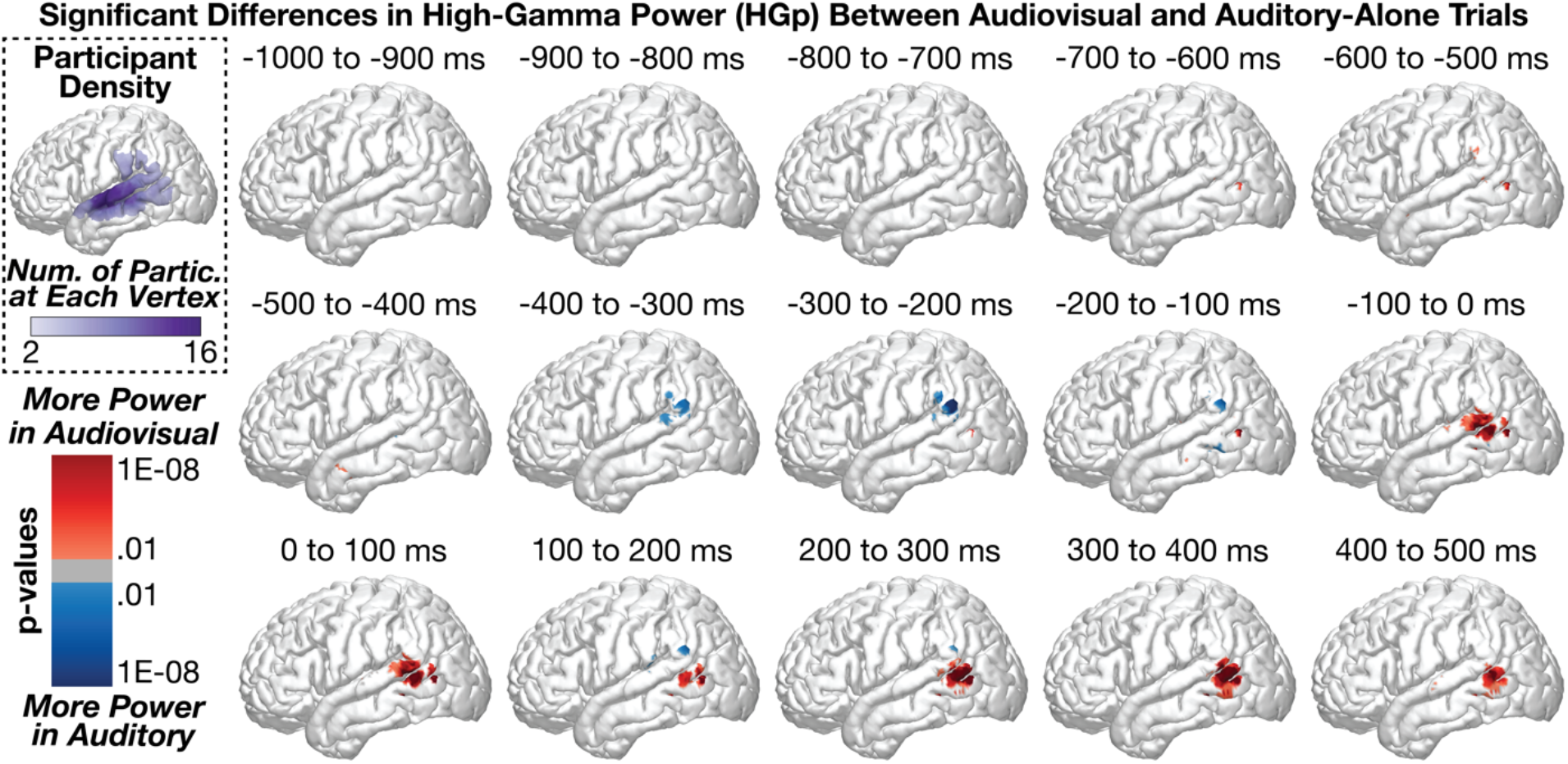
Group-level analyses comparing high gamma power (HGp) between audiovisual and auditory-alone conditions at 100 ms time-windows (sound onset a 0 ms). Top-left plot shows the number of participants who contributed data to each vertex. audiovisual stimuli elicited greater power at the posterior STG, peaking beginning before sound onset.

### 3.5 Group-Level Regional Time-Series Analyses

While the spatial analyses demonstrated significant patterns of activity along the STG, MTG, and SMG, this approach does not effectively allow comparisons across regions or allow the examination of interactions with time and across frequency. To model the influence of visual speech information on spectral power at the group level, we used linear mixed effects models for data aggregated into three regions of the STG (anterior, middle, and posterior regions), consistent with prior studies (Smith et al., 2013). Separate models were constructed at each time point and ROI, and multiple comparison corrections were applied. Importantly, in our estimation of condition effect (auditory-alone versus audiovisual), we modelled both random intercepts and slopes (Barr et al., 2013). Table 1 shows the number of electrodes and participants who contributed data to each analysis. Supplementary Figure 2 shows time-series analyses separated by task variants.

### 3.6 Group-Level Regional Time-Series Analyses: Theta Power

Regardless of condition, theta power within the STG increased steadily beginning before sound onset and peaking immediately after sound onset, with the strongest activity observed at the posterior STG. Consistent with the spatial analyses, we observed significant differences between audiovisual and auditory-alone conditions, with audiovisual trials demonstrating reduced auditory-related theta power (Figure 6). This condition difference was clearest at the posterior STG, which was significant from −93 to 500 ms (min *p* = 2.2e-06, peak time = 47 ms), yet also present at the middle STG, which was significant from 108 to 274 ms (min *p* = 0.030, peak time = 193 ms). No significant differences were observed at the anterior STG after correcting for multiple comparisons. To examine whether visual speech information differentially affected the three STG regions, we conducted a group-level linear mixed-effects model with additional factors of Time and ROI (see Methods for additional information). As expected, the effect of visual information varied as a function of time (Condition x Time interaction: [*F*(1, 2.0547e+06) = 851.6, *p* = 3.6e-187]), STG region (Condition x ROI interaction: [*F*(2, 2.0547e+06) = 28.7, *p* = 3.5e-13]) as well as the combination of the two (Condition x Time x ROI interaction: [*F*(2, 2.0547e+06) = 147.5, *p* = 8.8e-65]). The model with interaction terms additionally demonstrated better fit (AIC = 7.2815e+05) compared to the same model without interaction terms (AIC = 7.4294e+05). Taken together, these results indicate that visual speech information modulates auditory theta activity predominantly along the posterior STG, following sound onset.

**Figure 6.**
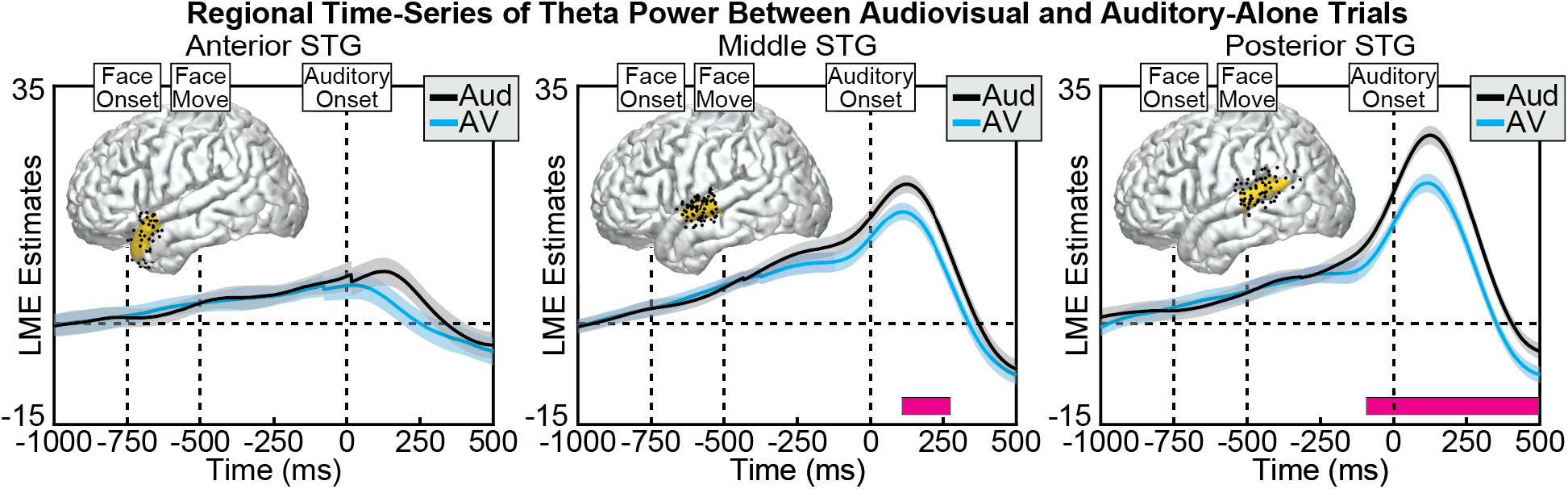
Group linear mixed-effect model (LME) estimates for each time-point of theta power in auditory-alone (black) and audiovisual (blue) trials, calculated separately at anterior (left), middle (middle), and posterior (right) regions of the STG. Shaded areas reflect 95% confidence intervals. Pink boxes reflect significant differences after correcting for multiple comparisons. Corresponding regions are highlighted on the cortical surfaces in yellow with the electrodes that contributed to the analysis shown as black dots (some depth electrodes are located beneath the surface and are not visible). Significant differences in theta power emerged largely after auditory, concentrated along the posterior STG.

### 3.7 Group-Level Regional Time-Series Analyses: Beta Power

Beta power in the STG showed a combination of power increases and power decreases (beta suppression), with the majority of activity focused on the mid- to posterior-STG. Across conditions, we observed significantly greater beta suppression during the audiovisual condition compared to the auditory-alone condition, peaking before sound onset at mid- to anterior STG regions (Figure 7). This condition difference was significant at both the anterior STG, significant from −311 to −195 ms (min *p* = 0.002, peak time = −247 ms), and the middle STG, significant from −195 to 235 ms (min p = 0.003, peak time = −116 ms), with no significant differences observed at the posterior STG after correcting for multiple comparisons. To examine whether visual speech information differentially affected the three STG regions, we conducted a group-level linear mixed-effects model with additional factors of Time and ROI. As expected, the effect of visual information varied as a function of time (Condition x Time interaction: [*F*(1, 1.9693e+06) = 48.5, *p* = 3.3e-12]), STG region (Condition x ROI interaction: [*F*(2, 1.9693e+06) = 44.2, *p* = 6.3e-20]) but a non-significant combination of the two (Condition x Time x ROI interaction: [*F*(2, 1.9693e+06) = 2.01, *p* = 0.134]). The model with interaction terms nevertheless demonstrated better fit (AIC = −1.3239e+05) compared to the same model without interaction terms (AIC = −1.2912e+05).

**Figure 7.**
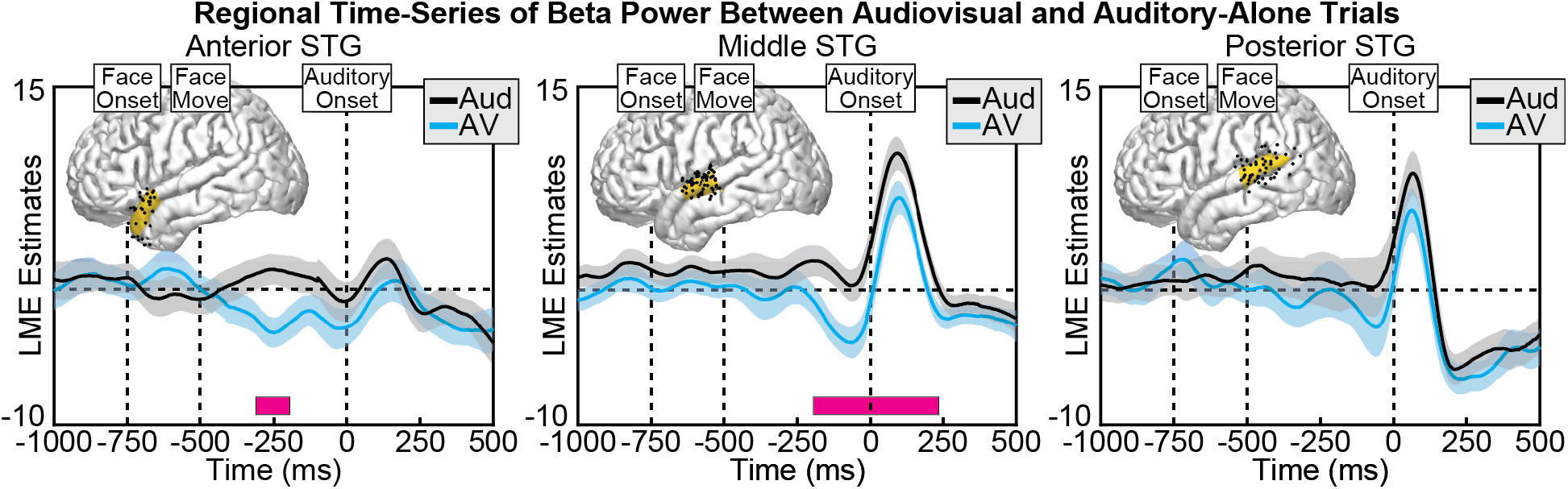
Group LME model estimates for each time-point of beta power in auditory-alone (black) and audiovisual (blue) trials, calculated separately at anterior (left), middle (middle), and posterior (right) regions of the STG. Pink boxes reflect significant differences after correcting for multiple comparisons. Corresponding regions are highlighted on the cortical surfaces in yellow with the electrodes that contributed to the analysis shown as black dots. Significant differences in beta power peaked before sound onset, concentrated in the middle to posterior STG.

### 3.8 Group-Level Regional Time-Series Analyses: High-Gamma Power

In general, HGp in the STG showed auditory-related power increases that were biased towards the posterior STG. Across conditions, we observed significantly greater HGp in the audiovisual condition compared to the auditory-alone condition, occurring before sound onset and localized to the posterior STG (Figure 8). This condition difference was significant only at the posterior STG, from −45 to 24 ms (min *p* = 0.028, peak time = −9 ms). No other significant differences were observed. To examine whether visual speech information differentially affected the three STG regions we conducted a group-level linear mixed-effects model with additional factors of Time and ROI. As expected, the effect of visual information varied as a function of time (Condition x Time interaction: [*F*(1, 1.7919e+06) = 86.7, *p* = 1.3e-20]), STG region (Condition x ROI interaction: [*F*(2, 1.7919e+06) = 29.6, *p* = 1.3e-13]) as well as the combination of the two (Condition x Time x ROI interaction: [*F*(2, 1.7919e+06) = 20.0, *p* = 2.1e-09]). The model with interaction terms additionally demonstrated better fit (AIC = −1.4021e+06) compared to the same model without interaction terms (AIC = −1.1389e+06).

**Figure 8.**
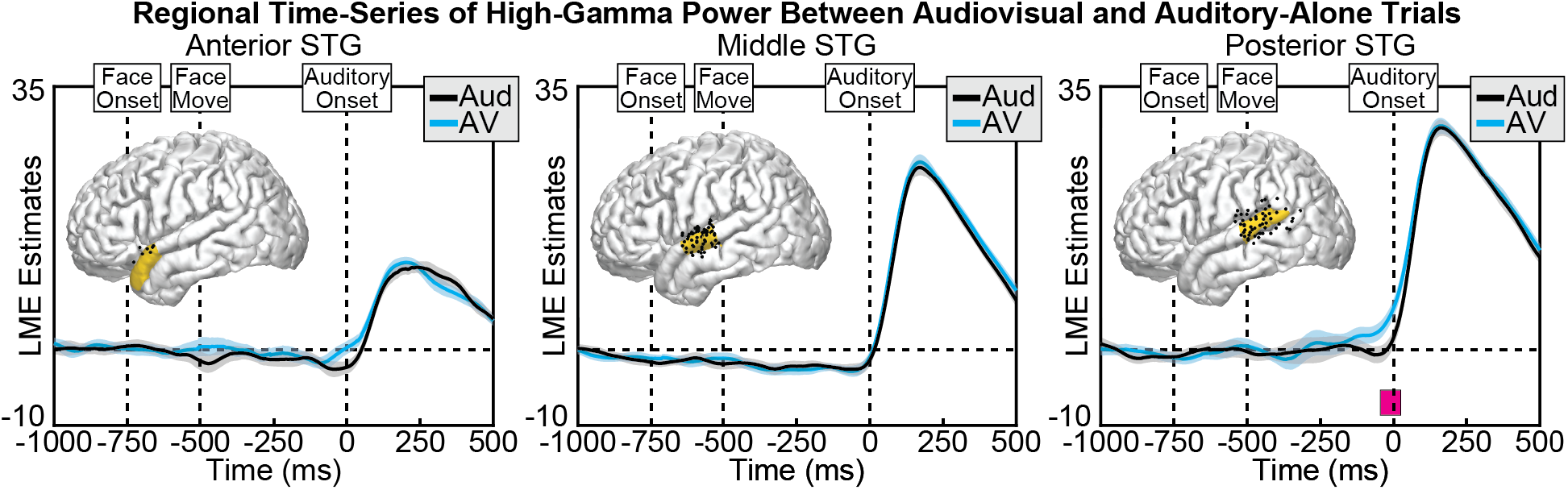
Group LME model estimates for each time-point of HGp in auditory-alone (black) and audiovisual (blue) trials, calculated separately at anterior (left), middle (middle), and posterior (right) regions of the STG. Pink boxes reflect significant differences after correcting for multiple comparisons. Corresponding regions are highlighted on the cortical surfaces in yellow with the electrodes that contributed to the analysis shown as black dots. Significant differences in HGp peaked before sound onset in the posterior STG.

### 3.9 Group-Level Regional Time-Series Analyses: Interactions Across Frequencies

Analyses conducted separately at each of the frequency bands demonstrated audiovisual effects in putatively distinct time ranges and spatial distributions. However, to test the claim that the spatial and temporal patterns observed across the frequency bands are indeed distinct, it is necessary to model frequency band and time-points in relation to task conditions. To this end, we constructed a group-level linear mixed-effects model that included fixed effects of task condition, frequency band, region of interest along the STG, and time, modeling both random intercepts and random slopes for trial condition. Including all frequency bands in the model yielded significant interactions of Condition x Frequency Band [*F*(2, 2.9e+07) = 277.4, *p* = 2.3e-64], Condition x Frequency Band x ROI [*F*(4, 2.9e+07) = 72.7, *p* = 4.8e-43], Condition x Frequency Band x Time [*F*(2, 2.9e+07) = 2254.0, *p* = 1.2e-294], and Condition x Frequency Band x ROI x Time [*F*(4, 2.9e+07) = 397.8, *p* = 3.7e-163]. Consistent with these significant interactions, the addition of each parameter improved model fit based on AIC. Repeating this analysis with only low-frequency signals associated with neural oscillations (theta and beta) yielded the same pattern, with significant interactions of Condition x Frequency Band [*F*(2, 2.0e+07) = 346.7, *p* = 2.2e-40], Condition x Frequency Band x ROI [*F*(2, 2.0e+07) = 43.2, *p* = 1.9e-15], Condition x Frequency Band x Time [*F*(1, 2.0e+07) = 1645.2, *p* = 2.5e-121], and Condition x Frequency Band x ROI x Time [*F*(2, 2.0e+07) = 357.6, *p* = 9.6e-78]. Again, the addition of each parameter improved model fit based on AIC. Taken together, these data demonstrate that visual speech information evokes distinct temporal and spatial patterns through theta, beta, and HGp.

### 3.10 Individual Differences in Neural Activity

While the linear mixed-effects models demonstrate effects that are present at the group level, it is important to note that highly significant condition differences that *deviated* from these group patterns were observed at individual electrodes in individual participants. In particular, HGp results showed greater variability across electrodes and participants than did theta and beta bands. For example, while the most consistently observed response was increased activity before sound onset in posterior regions of the STG, this was not present in all participants or all electrodes. Figure 9 shows pairs of individual electrode responses from 5 participants, with the top row highlighting one STG electrode from that participant that matches the pattern observed at the group level, and the bottom row highlighting a second STG electrode demonstrating a different (sometimes opposite) pattern. Indeed, Participant 9 (first column) showed the opposite pattern across two electrodes, with the lower row demonstrating more HGp for *auditory* trials before sound onset. Of note, many of the electrodes showed significantly reduced HGp to audiovisual versus auditory-alone stimuli during sound processing (100 - 200 ms), as reported previously (Karas et al., 2019). While this pattern was demonstrated in many electrodes and participants, the anatomical region varied throughout the STG and the overall pattern did not reach significance at the group level.

**Figure 9.**
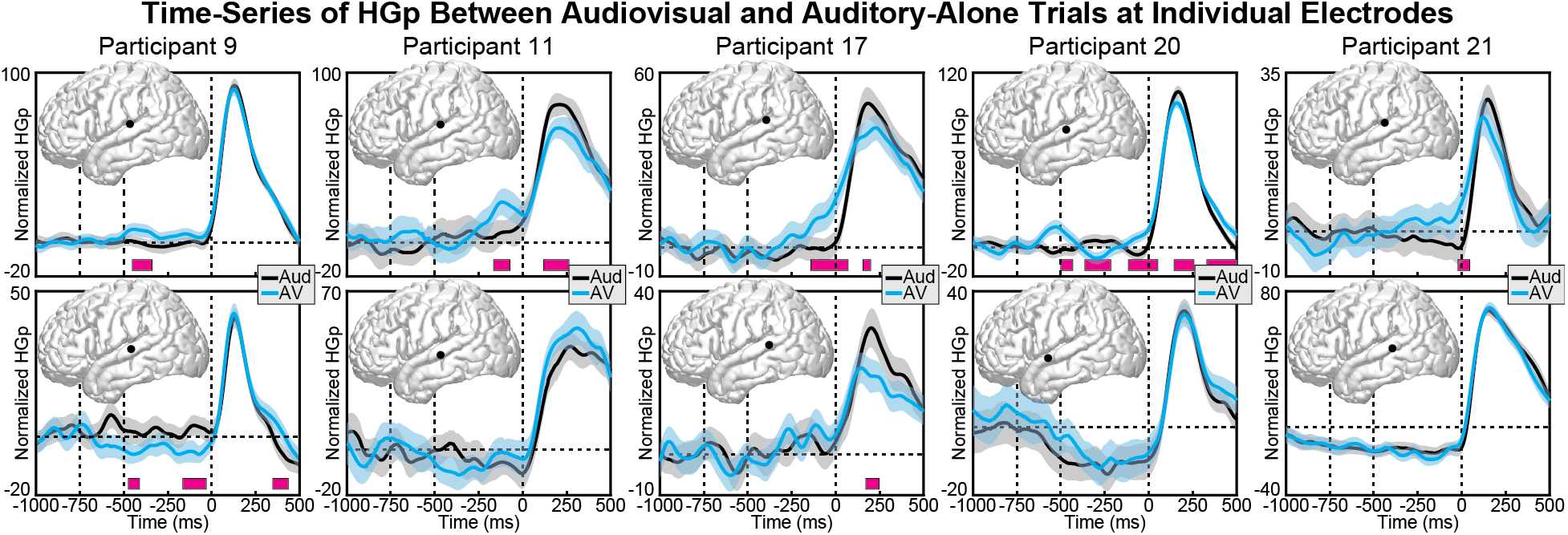
Individual participant HGp activity at audiovisual (blue) and auditory-alone (black) conditions. Each column displays data from a different participant (two electrodes per participant). Top row displays electrodes that showed the same pattern of HGp results observed at the group-level, with increased activity in the audiovisual condition starting before sound onset. Bottom row shows a proximal electrode that demonstrated a different (sometimes conflicting) pattern. Shaded areas reflect 95% confidence intervals. Pink boxes reflect significant differences after correcting for multiple comparisons.

## 4 Discussion

Visual signals are known to affect auditory speech processes in multiple ways. For example, lipreading signals provide high-level phonemic representations (Bourguignon et al., 2020), visual motion information can relay timing information (McGrath et al., 1985), lip closure facilitates the parsing of word-boundaries and speech rate (Chandrasekaran et al., 2009), lip-shape provides spectral information (Plass et al., 2020), and speaker identity can further enhance spatial localization and multisensory binding (Vatakis and Spence, 2007; Brang 2019). Indeed, a persistent challenge in identifying the various effects of audiovisual speech information has been largely methodological in nature. fMRI studies lack the temporal resolution to identify whether visual speech modulates auditory regions before, simultaneously with, or after the onset of auditory speech. On the other hand, iEEG studies face two critical shortcomings: (1) Past studies investigating audiovisual speech integration have analyzed data using single-participant designs or traditional parametric statistics making it hard to generalize the findings to the group-level and thus to the general population (Micheli et al., 2020; Besle et al., 2008; Plass et al., 2020). (2) Even while using variants of group-level analysis such as linear mixed-effects modeling, previous studies (Ozker et al., 2017; Ozker et al., 2018) have focused on HGp, which indexes local population firing rates, ignoring low-frequency oscillations which potentially reflect distinct audiovisual information.

To test for the presence of separate but concurrent visual processes in auditory areas, we measured neural activity using intracranially implanted electrodes in a large number of human participants (*n* = 21) during an audiovisual speech perception paradigm. These data demonstrated that at least three distinct patterns of activity occur in the STG during audiovisual speech perception relative to unimodal auditory speech perception. We observed (1) reduced beta power (higher beta suppression) in mid- to anterior-STG sites that peaked before sound onset, (2) increased high-gamma power localized to the posterior STG that also peaked before sound onset, and (3) reduced theta power in mid- to posterior-STG sites that peaked after sound onset. We interpret these distinct patterns to reflect distinct neural processing in auditory regions, potentially responsible for encoding different types of visual information to aid in auditory speech perception.

Converging behavioral and neurophysiological evidence suggests that audiovisual enhancements from audiovisual speech (e.g., better detection and faster reaction times) and visual recovery of phoneme information are subserved by two distinct mechanisms (Eskelund et al., 2011; Plass et al., 2014). This distinction may reflect a neural dissociation between predictive multisensory interactions that optimize feedforward encoding of auditory information and later feedback processes that alter auditory representations generated in the pSTS (Arnal et al., 2009; Arnal et al., 2011) and the posterior STG (Reale et al., 2007). In support of this view, both visual speech (Besle et al., 2004; Arnal et al., 2009; Van Wassenhove et al., 2005) and other anticipatory visual cues (Vroomen and Stekelenburg, 2010) can speed-up and reduce the magnitude of early physiological responses associated with auditory feedforward processing, potentially reflecting optimization of auditory encoding in accordance with temporal or acoustic constraints imposed by visual information. These early feedforward effects, which are insensitive to audiovisual congruity in speech, are temporally, spatially, and spectrally distinct from later (>300 ms) responses that are specific to crossmodally incongruent speech (Arnal et al., 2011; Van Wassenhove et al., 2005). These later incongruity-specific interactions point to a hierarchical feedback regime in which unisensory speech processing is altered in accordance with integrated audiovisual information from the pSTS (Olasagasti et al., 2015; Kayser and Logothetis, 2009) and general speech perception areas in the STG (Mesgarani et al., 2014). These data are consistent with this dissociation, with several temporally and spatially discrete neural responses in the STG. It should also be noted that some of these activation patterns may be due to non-specific effects (e.g., elevated attention or physiological arousal to viewing a face).

Our observation of a dissociation among theta and beta frequency ranges is consistent with prior EEG and physiology research suggesting these mechanisms encode different information about a visual signal (e.g., Kumar et al., 2016; Wang et al., 2017). Theta activity effectively captures ongoing auditory timing information, including rhythmic events (e.g., Schroeder et al., 2009). Conversely, beta band activity has been more strongly associated with feedback signals that may predictively encode visual information in the auditory system prior to sound onset (e.g., Engel et al., 2010). The dissociation between theta and HGp observed is particularly interesting as HGp signals have also been implicated in a predictive coding framework, such that ensembles of neurons in the posterior STG initially activate neuronal ensembles before sound onset, leading to refined population tuning and thus less HGp following sound onset (Karas et al., 2019). While this reduction in HGp during audiovisual trials was observed in many participants (see Figure 9), it was not observed at the group level, potentially due to anatomical variability in the location of the response or due to heterogeneity across participants.

Research on the neural source of visual signals relayed to the auditory system have largely focused on the left posterior temporal sulcus (pSTS). This region demonstrates strong differences between auditory-alone and audiovisual stimuli in both fMRI and iEEG research (Beauchamp et al., 2004b; Ozer et al., 2017; Ozker et al., 2018; Okada et al., 2013), and has potential causal roles in audiovisual speech integration as revealed by lesion mapping (Hickok et al., 2018) and inhibitory transcranial magnetic stimulation (Beauchamp et al., 2010). While these data indicate that some of the information observed in the present study was likely projected through feedback pathways originating in the pSTS, particularly given its role as a center for bottom-up prediction errors in language comprehension (Lewis and Bastiaansen, 2015), it is possible that each distinct temporal/spatial pattern has a unique corresponding source. While the present study does not provide evidence as to what information is encoded within each spatial/temporal pattern, we suggest that future research using causal measures or neural decoding identify the specific visual dimensions represented.

In summary, this study demonstrates that audiovisual speech integration elicits multiple distinct patterns of neural activity within the STG and adjacent cortex, occurring across separate frequencies and temporal/spatial distributions. These data suggest that visual modulation of auditory speech perception utilizes multiple mechanisms, potentially reflecting independent sources of information. Our results are also consistent a hybrid family of integration models as proposed by Peelle and Sommers (2015). Finally, this study additionally shows the advantage of group-level analyses of iEEG data using linear mixed-effect models, which can improve statistical validity and power, and importantly, improve generalization of results across patients and to the population at large.

## Acknowledgements

This study was supported by NIH Grant R00 DC013828 A. Beltz was supported by the Jacobs Foundation. The authors report no conflicts of interest.

## Supplementary Figures

**Supplementary Figure 1.**
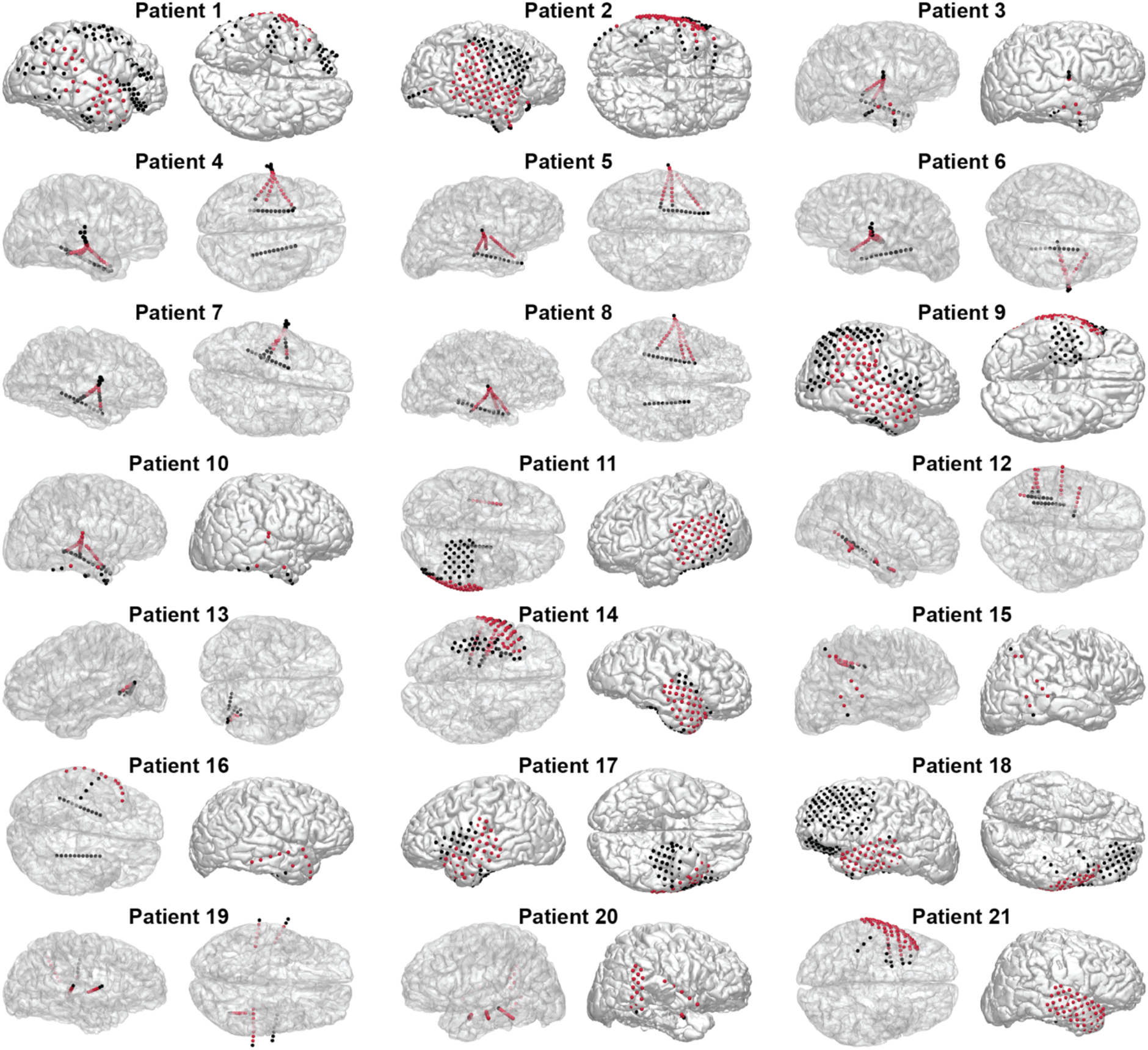
Supplementary Individual electrode maps for each patient. Red spheres reflect auditory electrodes that met anatomical criteria and that were not rejected during pre-processing for having excessive noise.

**Supplementary Figure 2.**
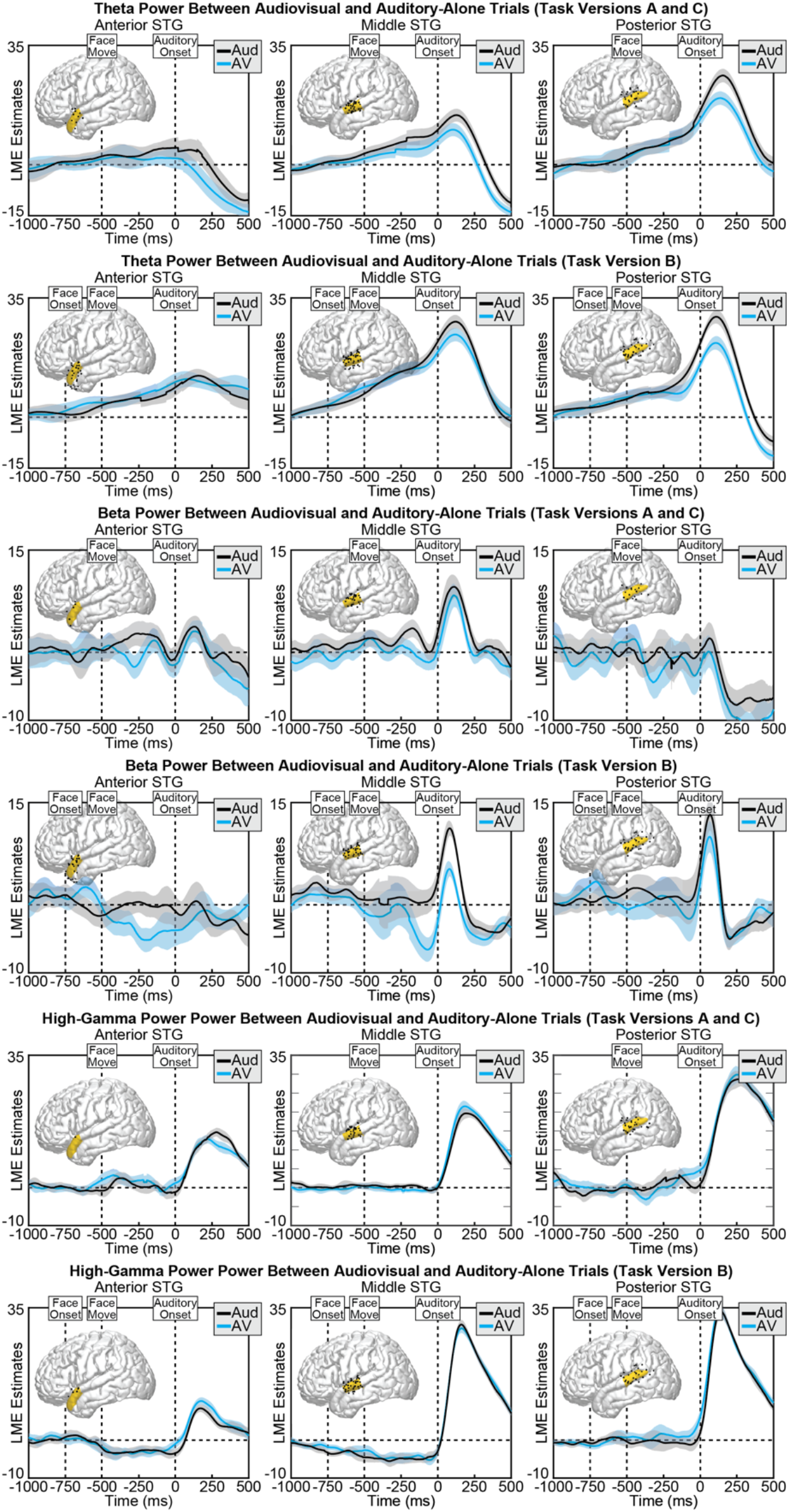
Individual participant activity at audiovisual (blue) and auditory-alone (black) conditions separated by task variants and frequency bands. Task Variants A and C presented participants the moving face stimulus at 500 ms before auditory onset, whereas Task Variant B showed a static face beginning 750 ms before auditory onset (with motion starting at 500 ms before auditory onset). No clear differences across the task differences are present in the time-series data.

## References

Aarts, E., Verhage, M., Veenvliet, J. V., Dolan, C. V., and Van Der Sluis, S. (2014). A solution to dependency: using multilevel analysis to accommodate nested data. Nature neuroscience, 17(4), 491–496.

Arnal, L. H., Morillon, B., Kell, C. A., and Giraud, A. L. (2009). Dual neural routing of visual facilitation in speech processing. Journal of Neuroscience, 29(43), 13445–13453.

Arnal, L. H., Wyart, V., and Giraud, A. L. (2011). Transitions in neural oscillations reflect prediction errors generated in audiovisual speech. Nature neuroscience, 14(6), 797.

Bastos, A. M., Usrey, W. M., Adams, R. A., Mangun, G. R., Fries, P., and Friston, K. J. (2012). Canonical microcircuits for predictive coding. Neuron, 76(4), 695–711.

Barr, D. J., Levy, R., Scheepers, C., & Tily, H. J. (2013). Random effects structure for confirmatory hypothesis testing: Keep it maximal. Journal of memory and language, 68(3), 255–278.

Beauchamp, M. S., Lee, K. E., Argall, B. D., and Martin, A. (2004a). Integration of auditory and visual information about objects in superior temporal sulcus. Neuron, 41(5), 809–823.

Beauchamp, M. S., Argall, B. D., Bodurka, J., Duyn, J. H., and Martin, A. (2004b). Unraveling multisensory integration: patchy organization within human STS multisensory cortex. Nature neuroscience, 7(11), 1190.

Beauchamp, M. S., Nath, A. R., and Pasalar, S. (2010). fMRI-Guided transcranial magnetic stimulation reveals that the superior temporal sulcus is a cortical locus of the McGurk effect. Journal of Neuroscience, 30(7), 2414–2417.

Beauchamp, M. S. (2016). Audiovisual speech integration: Neural substrates and behavior. In Neurobiology of language (pp. 515–526). Academic Press.

Bernstein, L. E., & Liebenthal, E. (2014). Neural pathways for visual speech perception. Frontiers in neuroscience, 8, 386.

Besle, J., Fort, A., Delpuech, C., and Giard, M. H. (2004). Bimodal speech: early suppressive visual effects in human auditory cortex. European journal of Neuroscience, 20(8), 2225–2234.

Besle, et al., (2008). Visual activation and audiovisual interactions in the auditory cortex during speech perception: intracranial recordings in humans. Journal of Neuroscience, 28(52), 14301–14310.

Bourguignon, M., Baart, M., Kapnoula, E. C., and Molinaro, N. (2020). Lip-reading enables the brain to synthesize auditory features of unknown silent speech. Journal of Neuroscience, 40(5), 1053–1065.

Brang, D., Dai, Z., Zheng, W., and Towle, V. L. (2016). Registering imaged ECoG electrodes to human cortex: A geometry-based technique. Journal of neuroscience methods, 273, 64–73.

Brang, D. (2019). The Stolen Voice Illusion. Perception, 48(8), 649–667.

Chandrasekaran, C., Trubanova, A., Stillittano, S., Caplier, A., and Ghazanfar, A. A. (2009). The natural statistics of audiovisual speech. PLoS computational biology, 5(7), e1000436.

Chang, E. F., Rieger, J. W., Johnson, K., Berger, M. S., Barbaro, N. M., and Knight, R. T. (2010). Categorical speech representation in human superior temporal gyrus. Nature neuroscience, 13(11), 1428.

Chen, T., and Rao, R. R. (1998). Audio-visual integration in multimodal communication. Proceedings of the IEEE, 86(5), 837–852.

Dale, A. M., Fischl, B., and Sereno, M. I. (1999). Cortical surface-based analysis: I. Segmentation and surface reconstruction. Neuroimage, 9(2), 179–194.

Elliott, T. M., and Theunissen, F. E. (2009). The modulation transfer function for speech intelligibility. PLoS comput biol, 5(3), e1000302.

Erber, N. P. (1975). Auditory-visual perception of speech. Journal of speech and hearing disorders, 40(4), 481–492.

Fischl, B., Sereno, M. I., and Dale, A. M. (1999). Cortical surface-based analysis: II: inflation, flattening, and a surface-based coordinate system. Neuroimage, 9(2), 195–207.

Hickok, G., Rogalsky, C., Matchin, W., Basilakos, A., Cai, J., Pillay, S., and Binder, J. (2018). Neural networks supporting audiovisual integration for speech: A large-scale lesion study. Cortex, 103, 360–371.

Engel, A. K., and Fries, P. (2010). Beta-band oscillations—signalling the status quo?. Current opinion in neurobiology, 20(2), 156–165.

Eskelund, K., Tuomainen, J., and Andersen, T. S. (2011). Multistage audiovisual integration of speech: Dissociating identification and detection. Experimental Brain Research, 208(3), 447–457.

Groppe, D. M., Urbach, T. P., and Kutas, M. (2011). Mass univariate analysis of event-related brain potentials/fields II: Simulation studies. Psychophysiology, 48(12), 1726–1737.

Hullett, P. W., Hamilton, L. S., Mesgarani, N., Schreiner, C. E., and Chang, E. F. (2016). Human superior temporal gyrus organization of spectrotemporal modulation tuning derived from speech stimuli. Journal of Neuroscience, 36(6), 2014–2026.

Kadipasaoglu, C. M., Baboyan, V. G., Conner, C. R., Chen, G., Saad, Z. S., and Tandon, N. (2014). Surface-based mixed effects multilevel analysis of grouped human electrocorticography. Neuroimage, 101, 215–224.

Kadipasaoglu, C. M., Forseth, K., Whaley, M., Conner, C. R., Rollo, M. J., Baboyan, V. G., and Tandon, N. (2015). Development of grouped icEEG for the study of cognitive processing. Frontiers in psychology, 6, 1008.

Kaiser, J., Hertrich, I., Ackermann, H., Mathiak, K., & Lutzenberger, W. (2005). Hearing lips: gamma-band activity during audiovisual speech perception. Cerebral Cortex, 15(5), 646–653.

Kaiser, J., Hertrich, I., Ackermann, H., & Lutzenberger, W. (2006). Gamma-band activity over early sensory areas predicts detection of changes in audiovisual speech stimuli. Neuroimage, 30(4), 1376–1382.

Karas, P. J., Magnotti, J. F., Metzger, B. A., Zhu, L. L., Smith, K. B., Yoshor, D., and Beauchamp, M. S. (2019). The visual speech head start improves perception and reduces superior temporal cortex responses to auditory speech. Elife, 8, e48116.

Kayser, C., and Logothetis, N. K. (2009). Directed interactions between auditory and superior temporal cortices and their role in sensory integration. Frontiers in integrative neuroscience, 3, 7.

Kleiner M, Brainard D, Pelli D, 2007, “What’s new in Psychtoolbox-3?” Perception 36 ECVP Abstract Supplement

Kumar, G. V., Halder, T., Jaiswal, A. K., Mukherjee, A., Roy, D., and Banerjee, A. (2016). Large scale functional brain networks underlying temporal integration of audio-visual speech perception: An EEG study. Frontiers in psychology, 7, 1558.

Lazic, S. E. (2010). The problem of pseudoreplication in neuroscientific studies: is it affecting your analysis?. BMC neuroscience, 11(1), 5.

Lazic, S. E., Clarke-Williams, C. J., and Munafò, M. R. (2018). What exactly is ‘N’in cell culture and animal experiments?. PLoS Biology, 16(4), e2005282.

Lega, B., Germi, J., and Rugg, M. D. (2017). Modulation of oscillatory power and connectivity in the human posterior cingulate cortex supports the encoding and retrieval of episodic memories. Journal of Cognitive Neuroscience, 29(8), 1415–1432.

Lewis, A. G., and Bastiaansen, M. (2015). A predictive coding framework for rapid neural dynamics during sentence-level language comprehension. Cortex, 68, 155–168.

Luke, S. G. (2017). Evaluating significance in linear mixed-effects models in R. Behavior research methods, 49(4), 1494–1502.

Mesgarani, N., Cheung, C., Johnson, K., and Chang, E. F. (2014). Phonetic feature encoding in human superior temporal gyrus. Science, 343(6174), 1006–1010.

McGrath, M., and Summerfield, Q. (1985). Intermodal timing relations and audio-visual speech recognition by normal-hearing adults. The Journal of the Acoustical Society of America, 77(2), 678–685.

Micheli, C., Schepers, I. M., Ozker, M., Yoshor, D., Beauchamp, M. S., and Rieger, J. W. (2020). Electrocorticography reveals continuous auditory and visual speech tracking in temporal and occipital cortex. European Journal of Neuroscience, 51(5), 1364–1376.

Okada, K., Venezia, J. H., Matchin, W., Saberi, K., and Hickok, G. (2013). An fMRI study of audiovisual speech perception reveals multisensory interactions in auditory cortex. PloS one, 8(6), e68959.

Ozker, M., Schepers, I. M., Magnotti, J. F., Yoshor, D., and Beauchamp, M. S. (2017). A double dissociation between anterior and posterior superior temporal gyrus for processing audiovisual speech demonstrated by electrocorticography. Journal of cognitive neuroscience, 29(6), 1044–1060.

Ozker, M., Yoshor, D., and Beauchamp, M. S. (2018). Converging evidence from electrocorticography and BOLD fMRI for a sharp functional boundary in superior temporal gyrus related to multisensory speech processing. Frontiers in human neuroscience, 12, 141.

Olasagasti, I., Bouton, S., and Giraud, A. L. (2015). Prediction across sensory modalities: A neurocomputational model of the McGurk effect. Cortex, 68, 61–75.

Peelle, J. E., and Sommers, M. S. (2015). Prediction and constraint in audiovisual speech perception. Cortex, 68, 169–181.

Plass, J., Guzman-Martinez, E., Ortega, L., Grabowecky, M., and Suzuki, S. (2014). Lip reading without awareness. Psychological science, 25(9), 1835–1837.

Plass, J., Brang, D., Suzuki, S., and Grabowecky, M. (2020). Vision perceptually restores auditory spectral dynamics in speech. Proceedings of the National Academy of Sciences, 117(29), 16920–16927.

Reale, R. A., Calvert, G. A., Thesen, T., Jenison, R. L., Kawasaki, H., Oya, H., … and Brugge, J. F. (2007). Auditory-visual processing represented in the human superior temporal gyrus. Neuroscience, 145(1), 162–184.

Riha, C., Güntensperger, D., Kleinjung, T., & Meyer, M. (2020). Accounting for Heterogeneity: Mixed-Effects Models in Resting-State EEG Data in a Sample of Tinnitus Sufferers. Brain topography, 33(4), 413–424.

Reuter, M., Rosas, H. D., & Fischl, B. (2010). Highly accurate inverse consistent registration: a robust approach. Neuroimage, 53(4), 1181–1196.

Schroeder, C. E., Lakatos, P., Kajikawa, Y., Partan, S., and Puce, A. (2008). Neuronal oscillations and visual amplification of speech. Trends in cognitive sciences, 12(3), 106–113.

Schroeder, C. E., and Lakatos, P. (2009). Low-frequency neuronal oscillations as instruments of sensory selection. Trends in neurosciences, 32(1), 9–18.

Smith, E., Duede, S., Hanrahan, S., Davis, T., House, P., and Greger, B. (2013). Seeing is believing: neural representations of visual stimuli in human auditory cortex correlate with illusory auditory perceptions. PLoS One, 8(9), e73148.

Stouffer, S. A., Suchman, E. A., DeVinney, L. C., Star, S. A., and Williams, R. M., Jr. (1949). The American Soldier: Adjustment During Army Life (Vol. 1). Princeton, NJ: Princeton University Press.

Van Wassenhove, V., Grant, K. W., and Poeppel, D. (2005). Visual speech speeds up the neural processing of auditory speech. Proceedings of the National Academy of Sciences, 102(4), 1181–1186.

Vatakis, A., and Spence, C. (2007). Crossmodal binding: Evaluating the “unity assumption” using audiovisual speech stimuli. Perception and psychophysics, 69(5), 744–756.

Vroomen, J., and Stekelenburg, J. J. (2010). Visual anticipatory information modulates multisensory interactions of artificial audiovisual stimuli. Journal of Cognitive Neuroscience, 22(7), 1583–1596.

Wang, L., Wang, W., Yan, T., Song, J., Yang, W., Wang, B., … and Wu, J. (2017). Beta-band functional connectivity influences audiovisual integration in older age: an EEG study. Frontiers in aging neuroscience, 9, 239.

Wang, X. J. (2010). Neurophysiological and computational principles of cortical rhythms in cognition. Physiological reviews, 90(3), 1195–1268.

